# BitMapperBS: a fast and accurate read aligner for whole-genome bisulfite sequencing

**DOI:** 10.1101/442798

**Authors:** Haoyu Cheng, Yun Xu

## Abstract

As a gold-standard technique for DNA methylation analysis, whole-genome bisulfite sequencing (WGBS) helps researchers to study the genome-wide DNA methylation at single-base resolution. However, aligning WGBS reads to the large reference genome is a major computational bottleneck in DNA methylation analysis projects. Although several WGBS aligners have been developed in recent years, it is difficult for them to efficiently process the ever-increasing bisulfite sequencing data. Here we propose BitMapperBS, an ultrafast and memory-efficient aligner that is designed for WGBS reads. To improve the performance of BitMapperBS, we propose various strategies specifically for the challenges that are unique to the WGBS aligners, which are ignored in most existing methods. Our experiments on real and simulated datasets show that BitMapperBS is one order of magnitude faster than the state-of-the-art WGBS aligners, while achieves similar or better sensitivity and precision. BitMapperBS is freely available at https://github.com/chhylp123/BitMapperBS.

## 1 Introduction

DNA methylation plays important roles in many biological processes, such as genomic imprinting, silencing of transposable elements, cell proliferation and differentiation [1]. It has been proven to be associated with a number of human diseases, especially with cancer [2, 3, 4]. Recent advance of whole-genome bisulfite sequencing (WGBS) makes it possible to unbiasedly profile the methylation status of the entire genome. To obtain WGBS reads, DNA is first treated with sodium bisulfite, and then sequenced by high-throughput sequencing technologies. As a result, the unmethylated cytosines (C) are converted to thymines (T), while the methylated Cs are still unchanged. For existing WGBS analysis, the fundamental step is to align massive WGBS reads to the reference genome. A specific problem of WGBS aligners, termed asymmetric match, is that they need to align Ts in reads to Cs in the reference genome, but not vice versa. Besides, compared with the DNA sequence reads, the WGBS reads present relatively lower sequence complexity as most Cs are converted to Ts. In practice, the low sequence complexity of WGBS reads may degrade the performance of short read aligners. For example, existing aligners usually adopt the well-known seed-and-extend strategy. This strategy first identifies the occurrence positions of seeds (short subsequences of reads) in the reference genome, and then extends these occurrence positions, called candidate locations, to obtain the full alignment of each read. Owning to the lower sequence complexity of WGBS reads, there are more occurrence positions of their seeds. Given a large reference genome, the WGBS aligners are time-consuming since they need to extend massive candidate locations.

To solve these problems, a number of WGBS aligners have been proposed. Generally, these methods can be classified into two types: wild-card aligners and three-letter aligners. Wild-card aligners including BSMAP [5] and RMAPBS [6], introduce a new wild-card to allow the asymmetric match between Cs and Ts. By contrast, recent aligners like Bismark [7], WALT [8], BRAT-nova [9], BS-Seeker2 [10], BS-Seeker3 [11] and GEMBS [27], adopt the three-letter strategy. This strategy first converts all Cs to Ts in both the reads and the reference genome, and then aligns the CT-converted reads to the CT-converted reference genome. Finally, the best mapping location of each read is identified from the alignment results based on the original read and the original reference genome. By utilizing the wild-card strategy or the three-letter strategy, existing WGBS aligners can support the asymmetric match between Cs and Ts. However, the underlying data structures and algorithms of existing WGBS aligners are similar to those of traditional DNA sequence aligners [12, 13], which cannot process the WGBS reads with low sequence complexity efficiently. With the explosive growth of WGBS data, there is an urgent need to develop more efficient WGBS aligners.

Here we present BitMapperBS, an ultra-fast and memory-efficient aligner that is designed for WGBS reads from directional protocol. In practice, almost all bisulfite sequencing reads are generated using directional protocol, which only sequences two original strands (i.e., the original forward strand and the original reverse complement strand). To support inexact alignment of WGBS reads, BitMapperBS follows the three-letter strategy and the seed-and-extend strategy. The high performance of BitMapperBS is mainly attributed to three factors. First, we propose a novel version of FM-index specifically for the three-letter bisulfite sequencing data. In contrast, the FM-indexes [14] used in existing WGBS aligners are similar to those in common DNA sequence aligners, which are optimized for the four-letter DNA sequence data instead of the three-letter data. For the FM-index, ranking over the Burrows Wheeler Transform (BWT) [15] is the basic and critical operation [16]. BitMapperBS reorganizes BWT to accelerate the rank operations when the alphabet size is three. Second, in extension step, BitMapperBS adopts the vectorized bit-vector algorithm [17] to calculate the full alignment efficiently. Compared with DNA sequence aligners, WGBS aligners need to extend more candidate locations due to the lower sequence complexity of bisulfite sequencing reads. The vectorized bit-vector algorithm used in BitMapperBS extends multiple candidate locations simultaneously, while existing aligners extend their candidate locations one-by-one. As a result, the time-consuming extension step of BitMapperBS can be significantly accelerated. We also modifiy this algorithm to directly support the asymmetric match between Cs and Ts. Third, we propose several other techniques specifically for the problems that are unique to the WGBS aligners. Based on the above factors, BitMapperBS is up to 70 times faster than popular WGBS aligners Bismark [7] and BSMAP [5], and presents similar or better level of sensitivity and precision. In addition, BitMapperBS is memory-efficient, so that it is able to align massive WGBS reads to large human genome on a standard desktop computer with 8 GB RAM.

## 2 Methods

BitMapperBS is designed to efficiently align massive WGBS reads against the large reference genome. It is based on the three-letter strategy and the seed-and-extend strategy. During the index building phase, BitMapperBS builds a highly optimized FM-index on the concatenation of the reference genome’s two original strands with all Cs converted to Ts. By utilizing the precomputed FM-index, BitMapperBS first searches for the occurrence positions of short seeds in seeding step. After that, in extension step, these occurrence positions of seeds, termed candidate locations, are extended to obtain the full alignment of each read. Like most existing WGBS aligners [5, 7, 8, 9, 10, 27], BitMapperBS discards ambiguous read if it is equally aligned to multiple best mapping locations. Owning to the low sequence complexity of WGBS reads, randomly reporting one or a few best mapping locations of an ambiguous read may lead to incorrect alignment result.

Although the WGBS aligners and the traditional DNA sequence aligners share same seed-and-extend strategy, there are a number of problems that are unique to the WGBS aligners. Actually, existing WGBS aligners mainly focus on the most obvious problem - that is, how to support the asymmetric match between Cs and Ts. However, surprisingly little attention has been devoted to several underlying problems, which may degrade the performance of WGBS aligners. Here we briefly describe each step of BitMapperBS, and present the strategies that are used to solve these underlying problems. The novel FM-index designed specifically for the three-letter bisulfite sequencing data is also discussed in the following sections.

### 2.1 The three-letter FM-index

As a compressed full-text index, the FM-index is proposed to reduce the memory footprint of the classical suffix array (SA) [18]. Given a reference genome *G*, its suffix array *SA*(*G*) saves the positions of all suffixes of *G* in lexicographic order. For a seed *S* which is searched via *SA*(*G*), its all occurrence positions are contiguously saved in an interval in *SA*(*G*). The reason is that all suffixes prefixed by *S* are sorted together. Once the SA interval of seed *S* is obtained, all occurrence positions of it can be directly retrieved. Formally, the operation that is used to determine the SA interval is termed as counting operation. To reduce the memory footprint of suffix array, the general idea of FM-index is to save a small fraction of positions of the reference genome, instead of all of them. We refer to these positions as sampled positions. Like suffix array, FM-index first performs counting operation when searching for seed *S* via it. However, another time-consuming operation, called locating operation, is also required for FM-index. The locating operation of FM-index is designed to recover the unsampled positions of *S*. We refer to [16, 19, 20] for more details of the counting operation and the locating operation of FM-index.

For the FM-index, both the counting operation and the locating operation are based on the LF-mapping operation [16, 19, 20], and finally can be reduced to rank operation over the Burrows Wheeler Transform (BWT) [15]. As the critical data structure of the FM-index, the BWT of a reference genome *G* is a permutation of *G*. Specifically, *BWT*[*i*] = *G*[*SA*[*i*] – 1] if *SA*[*i*] ≠ 0, otherwise *BWT*[*i*] = $, where $ is a sentinel character that is lexicographically smaller than all characters in *G*. The rank operation *rank_c_*(*BWT*; *i*) over BWT returns the number of character *c* in *BWT*[0, *i* – 1], where *BWT*[0, *i* – 1] is the *BWT*’s prefix of length *i*. We also let *interval_rank_c_*(*BWT, i, j*) = *rank_c_*(*BWT, j*) *rank_c_*(*BWT, i*). In practice, existing aligners exploit a precomputed auxiliary data structure *Occ* to support fast rank operation. In these aligners, BWT is organized into small blocks of equal length. For the *m*-th block of BWT and each character *c* of alphabet, *Occ*[*m, c*] = *rank_c_*(*BWT, m * b*), where *b* is the length of each block. An example of *Occ* is presented in Figure 1. When *rank_c_*(*BWT, i*) is required, one can calculate it as follows: *rank_c_*(*BWT, i*) = *Occ*[[*i/b, c*] + *interval_rank_c_*(*BWT*, [*i/b*] * *b,i*). Since *interval_rank_c_*(*BWT*, [*i/b*] * *b,i*) needs to be carried out on the fly, it is much more computationally expensive than retrieving *Occ*[[*i/b*, *c*].

**Figure 1.**
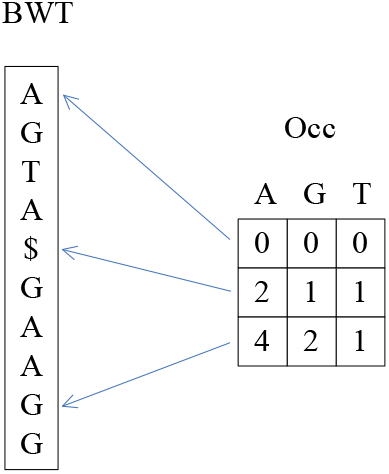
An example of *Occ* array used to accelerate the rank operation over BWT. In this example, BWT = “*AGTA$GAAGG*” consists of 10 characters, and the length of each block in BWT is 4. For three-letter WGBS aligners, the alphabet of BWT only includes *A*, *G* and *T*.

Essentially, *interval_rank_c_*(*BWT*, [*i/b*] * *b,i*) counts the number of character *c* in *BWT*[[*i/b*] * *b,i* – 1]. Although a number of attempts have been made to support this operation [19, 20], most existing aligners adopt a simple method for genomic data [21, 22, 23]. This method represents each character of BWT by two bits. That is, *A*, *C*, *G* and *T* are encoded as *code*(*A*) = 00, *code*(*C*) = 01, *code*(*G*) = 10 and *code*(*T*) = 11, respectively. Actually, the practical implementation of BWT is a 64-bit word (64-bit integer) array, where every element of it represents 32 characters (see Figure 2 (a)). As a result, *interval_rank_c_*(*BWT*, [*i/b*] * *b,i*) is reduced to the rank operation over a 64-bit word (i.e., counting the number of *code*(*c*) in a 64-bit word). Here we refer to this operation as character count operation, and let *c_count(c, w)* be the character count operation of character *c* over 64-bit word *w*. For existing aligners, there are two efficient strategies for this operation instead of scanning all characters one-by-one:

- **Lookup table based method:** For each character of alphabet, this method constructs a static lookup table in advance. When the number of character *c* in *m*-bit words is required, the size of its corresponding lookup table *lookup_c_* equals to *2*^m^. The *i*-th element *lookup_c_*[*i*] (0 ≤ i ≤ 2^m^—1) in *lookup_c_* stores the number of *code*(*c*) in *m*-bit word *i*. In most cases, 8-bit lookup tables or 16-bit lookup tables are used in existing aligners, since these small lookup tables can be kept in fast but small CPU cache. By utilizing the 16-bit lookup tables, the number of character *c* in a 64-bit word *w* is able to be calculated as follows: *c_count*(*c*, *w*) = *lookup_c_*[*w* ≫ 48] + *lookup_c_*[(*w* ≫ 32) & 65535] + *lookup_c_*[(*w* ≫ 16) & 65535] + *lookup_c_*[*w* & 65535].
- **Built-in function based method:** This method is developed based on a CPU built-in function *_builtin_popcountll*, which returns the number of bit 1 in a given 64-bit word. A practical implementation of this method is presented in Supplementary Section S1. Its general idea is to first set related bits of queried character *c* to 1, and set other bits to 0. After that, the number of *c* is calculated by identifying the number of bit 1 using *_builtin_popcountll*. In total, to perform character count operation over a 64-bit word representing 32 characters, this built-in function based method needs at least four bit operations and one built-in operation *_builtin_popcountll* (see Supplementary Section S1).

**Figure 2.**
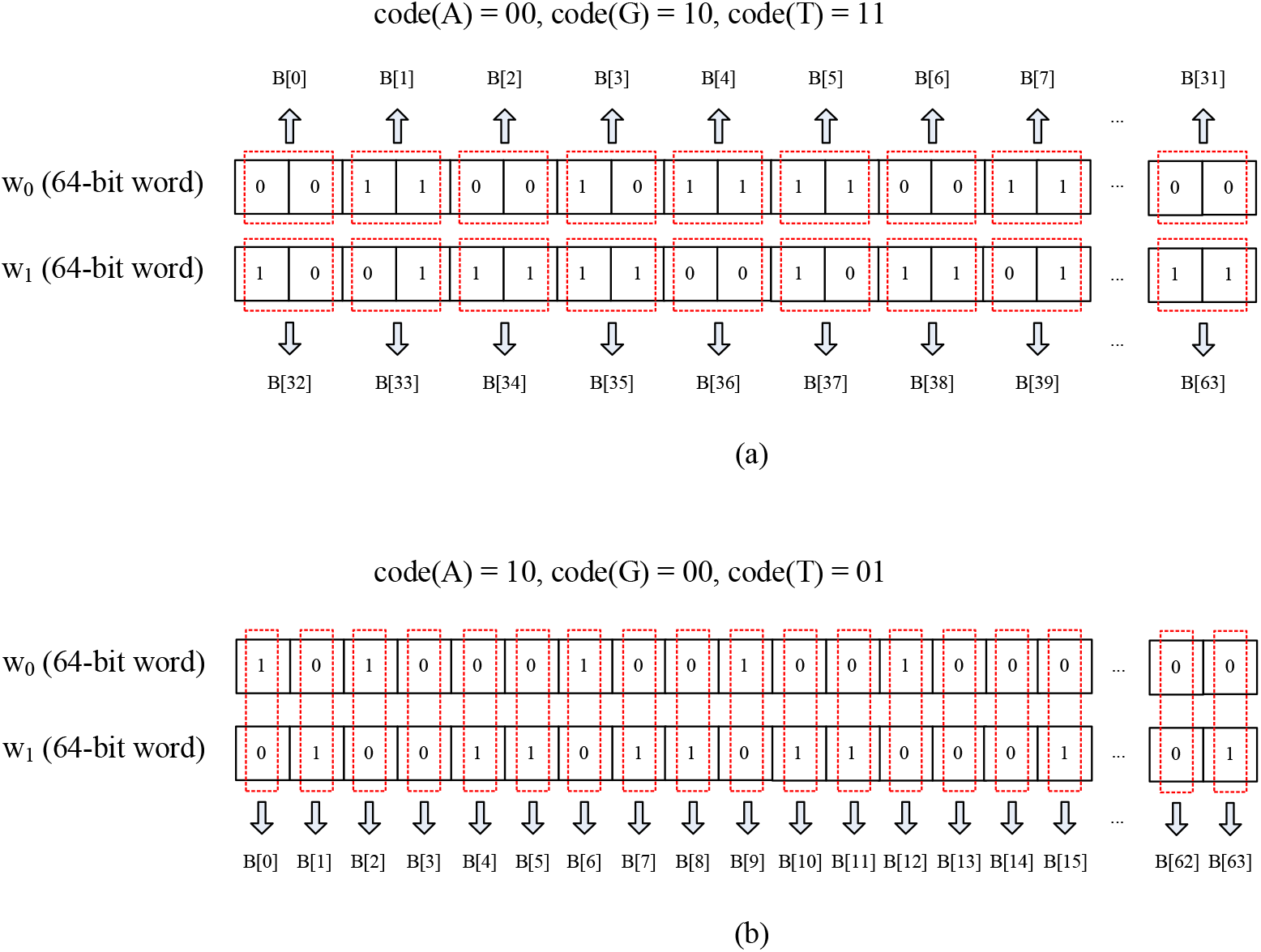
An example of practical data structures of BWT. For most existing aligners, BWT is represented by a 64-bit word array. In this example, the toy BWT sequence B = “*ATAGTTATTATTAGGT…GT*” consists of 64 characters, and two 64-bit words *w*_0_ and *w*_1_ are used to represent B. (a) The data structure of BWT in existing four-letter FM-index. (b) The data structure of BWT in our three-letter FM-index.

Although the above methods present excellent performance for four-letter DNA sequence data, there is no method that is designed specifically for three-letter bisulfite sequencing data. In BitMapperBS, we propose a three-letter FM-index which reorganizes BWT to accelerate the character count operation. Like existing implementations of FM-index, the three-letter FM-index encodes each character of BWT by two bits. In this case, *A, G* and *T* are encoded as *code*(*A*) = 10, *code*(*G*) = 00 and *code*(*T*) = 01, respectively. Note that *C* is not encoded by three-letter FM-index, since all Cs have been converted to Ts in BitMapperBS. To support fast character count operation for three-letter data, our three-letter FM-index stores two bits of 2-bit code of each BWT’s character in different 64-bit words. In contrast, for existing FM-index, all bits of the 2-bit code of each character in BWT are saved contiguously in a 64-bit word. An example of the BWT in our three-letter FM-index is presented in Figure 2 (b). Besides, another example of the BWT in existing four-letter FM-index can be found in Figure 2 (a). Specifically, let *c* be the *i*-th character in its corresponding block of BWT, and *BBWT* be the 64-bit word array of *c*’s corresponding block in BWT. We assume that *BBWT*[*j*][*k*] is the *k*-th bit in the *j*-th 64-bit word of *BBWT*. The three-letter FM-index stores the 0-th bit of *code*(*c*) in *BBWT*[[*i*/64] * *2*][*i* mod 64], and the *1*-th bit of *code*(*c*) in *BBWT*[[*i*/64] * 2 + 1][*i* mod 64]. In other words, all characters in BWT are organized into groups of 64. For every 64 characters, the 0-th bits of their 2-bit codes are saved in a 64-bit word, and the 1-th bits of their 2-bit codes are saved in another 64-bit word.

By utilizing the reorganized BWT, the character count operation for three-letter data can be significantly accelerated. To represent every 64 characters, the three-letter BWT needs two 64-bit words. Intuitively, the 0-th bits of these characters’ 2-bit codes are saved in the first 64-bit word *w*_0_, and the *1*-th bits of these characters’ codes are saved in the second 64-bit word *w*_1_. Recall that *code*(*A*) = 10, *code*(*G*) = 00 and *code*(*T*) = 01 in BitMapperBS. Obviously, the number of *A* among these 64 characters is able to be calculated using one built-in operation *_builtin_popcountll*(*w*_0_). This is because only the 0-th bit of *code*(*A*) is 1. Similarly, the number of *T* can be calculated by one operation *_builtin_popcountll*(*w*_1_). To obtain the number of *G* among 64 characters, we need two extra bit operations before *_builtin_popcountll*. First, a bit-XOR operation between *w*_0_ and *w*_1_ is required. Since two bits of *code*(*G*) are same, this operation produces a new 64-bit word that only the related bits of *G* are set to 0. Second, a bit-inversion operation is used to make the related bits of *G* be 1 and other bits be 0. As a result, the number of *G* can be determined by *_builtin_popcountll*. Formally, the character count operation for *G* is carried out as follows: *_builtin_popcountll*(∼(*w*_0_ ⊕ *w*_1_)). Therefore, given 64 characters, counting the numbers of *A*, *G* and *T* requires 1, 3 and 1 operations, respectively.

For three-letter WGBS aligners, the frequencies of both *A* and *G* in queried seeds are about 25%. However, the frequency of *T* is about 50%, since all Cs have been converted to Ts in the seeding step of these three-letter aligners. In our three-letter FM-index, the expected time of character count operation over 64 characters is 1∗ 25%+3∗25%+1∗ 50% = 1.5. As mentioned above, the existing methods based on traditional four-letter FM-index need at least 5 operations to count a character over 32 characters, so that at least 10 operations are required when counting a character over 64 characters. If we simply assume that the cost of different operations is equal, our method is at least 6.7 times faster than existing methods when processing three-letter data. In fact, the character count operation is the most computationally expensive part in rank operation, and rank operation over BWT is the basic and critical operation in searching algorithm of FM-index. Therefore, by utilizing our three-letter FM-index, BitMapperBS can search short seeds more efficiently during the seeding step.

### 2.2 Other optimized strategies for FM-index

In current WGBS aligners, there are two problems about the index that do not exist in traditional DNA sequence aligners. First, during the seeding step, three-letter WGBS aligners need to search for the seeds extracted from the CT-converted reads via the index built on the CT-converted reference genome. In this case, both the seeds and the reference genome with 3-letter alphabet present low sequence complexity. These seeds consist of much more occurrence positions than the seeds used in four-letter DNA sequence aligners. For the FM-index, a major bottleneck is that its locating operations are extremely slow when searching for short seeds with many occurrence positions [16]. Therefore, the seeding step in FM-index based WGBS aligners is more time-consuming than that in DNA sequence aligners. Second, to accelerate the WGBS alignment, recent aligners [9, 11, 27] build a single index on the concatenation of the forward strand and the reverse complement strand of the reference genome. For human genome, its concatenation of two strands consists of about 6.2 billion characters. However, the indexes used in existing aligners usually store each position of the reference genome by a 32-bit integer, which cannot handle large sequence (i.e., the concatenation of two strands) with more than 2^32^ — 1 ≈ 4.3 billion characters. To solve the first problem, existing aligners [8, 11] build uncompressed indexes for the reference genome to avoid the time-consuming locating operations. However, the use of uncompressed indexes would lead to large memory requirement of these aligners. For example, WALT builds four uncompressed hash table indexes for the human genome, each of which requires about 14 GB RAM. To solve the second problem, some WGBS aligners represent each position of the reference genome with more than 32 bits. This strategy not only increases the memory requirement, but also results in extra operations when retrieving positions from indexes.

In BitMapperBS, we adopt an efficient locating algorithm FMtree [16] to address the above problems. This algorithm recovers the unsampled positions of FM-index block-by-block, while the locating algorithm used in existing WGBS aligners recovers these positions one-by-one. In particular, FMtree is well suited for three-letter WGBS aligners, since it presents better performance for the patterns (i.e., seeds) with more occurrence positions and smaller alphabet size. The original implementation of FMtree is proposed for four-letter DNA sequence data. In BitMapperBS, we redesigned it for three-letter data to further improve its performance. Given a queried seed, we organize the search space of its locating operations into a ternary tree, and calculate its all unsampled positions by traversing this ternary tree. Instead, the search space of the original implementation of FMtree is a quadtree, which needs more time to be traversed. By utilizing the compressed FM-index and its fast locating algorithm FMtree, BitMapperBS is memory-efficient without sacrificing speed. For example, BitMapperBS required 6.8 GB RAM when aligning WGBS reads to the human genome, while the second fastest aligner GEMBS required more than 30 GB RAM (see Table 2 in Section 3). Moreover, BitMapperBS was about one order of magnitude faster than GEMBS in our experiments. Another advantage of FMtree is that it also solves the second problem mentioned above. To perform FMtree, the FM-index must be sampled by value sampling strategy. This strategy samples every *SA*[*i*] if *SA*[*i*] = 0 (mod *D*), where *D* is the sampling value. In practice, for each sampled position *SA*[*i*], *SA*[*i*]/*D* is saved in the FM-index instead of *SA*[*i*]. By default, the sampling value *D* of FM-index is set to 8 in BitMapperBS. When processing the concatenation of the human genome’s two strands including 6.2 billion characters, the maximum sampled position of FM-index is not more than 6.2 billion / 8 ≈ 775 million. As a result, it is enough for BitMapperBS to represent each sampled position of the FM-index by a 32-bit word, which is able to handle large sequence including at most 4.3 billion characters.

### 2.3 Seed-and-extend strategy

In seeding step, BitMapperBS extracts multiple overlapping and variable-length seeds from every read, and searches for the occurrence positions of these seeds exactly via the three-letter FM-index. Given a queried read *R*, each seed of it is a subsequence *R*[*i, j*] that starts at *R*[*i*] and ends at *R*[*j*]. By utilizing the FM-index, these seeds are aligned character-by-character until any of two following criteria are met. First, if the number of occurrence positions of *R*[*i, j*] is 1, *R*[*i, j*] is selected as a seed. Second, once there are no occurrence positions of *R*[*i, j*] in the reference genome, *R*[*i, j* — 1] is extracted as a seed. For different reads, we adopt a flexible strategy that selects different numbers of seeds. Its key idea is to extract less seeds for the reads that are easy to be aligned, and extract more seeds for the reads that are difficult to be aligned. Specifically, when a seed *R*[*i, j*] of length *n = j — i +* 1 has been extracted from read *R*, BitMapperBS selects next seed starting from *R*[*i* + [*n*/2]]. In contrast to the reads with many short seeds, the reads with longer seeds generate less seeds in BitMapperBS. Intuitively, if several long seeds of a read can be exactly aligned to the reference genome, there may be no or a few errors in this read. In this case, we select less seeds to reduce the running time without sacrificing alignment sensitivity. By contrast, a read without long seeds is more likely to include more errors. Thus, we need to select more seeds to avoid incorrect alignment. It is also noteworthy that in BitMapperBS, the exactly CT-converted seeds can be aligned with mismatches between Cs and Ts. For example, when a C in the original WGBS read is aligned to a *T* in the original reference genome, the three-letter strategy ignores this mismatch so that it is allowed in the exactly CT-converted seeds. This also suggests that selecting longer seeds may not sacrifice sensitivity of BitMapperBS. Actually, recent DNA and RNA sequence aligners [24, 25] present satisfactory sensitivities with even less seeds than BitMapperBS. This is because the sequencing errors in high quality reads produced by new sequencing technologies have been significantly reduced. And in practice, all reads need to be preprocessed before alignment by quality trimming. This operation further reduces the errors of reads.

In extension step, most existing aligners only extend a fraction of seeds’ occurrence positions that are more likely to be aligned. We refer to these positions as candidate locations. For four-letter DNA sequence aligners, this strategy is able to accelerate the extension step, and does not sacrifice the alignment sensitivity and precision. However, owning to the low sequence complexity of WGBS reads, it is difficult for WGBS aligners to identify the potential mapping locations without full alignment. This is also why most WGBS aligners only report the results of unique aligned reads. In this case, if only a fraction of seeds’ occurrence positions are extended, both the sensitivities and precisions of WGBS aligners may be degraded. To solve this problem, BitMapperBS selects all of the occurrence positions of seeds as candidate locations, and calculates the full alignment of them during extension step. As shown in Table 1 in Section 3, BitMapperBS presented the best precision among all WGBS aligners. This is reasonable, since most incorrect alignments are able to be discarded by the exhaustive extension strategy of BitMapperBS.

**Table 1.** Results of 1 million simulated paired-end 150bp reads from human genome.

| Aligner | No. of unique aligned reads | No. of correctly aligned reads | No. of incorrectly aligned reads | Sensitivity (%) | Precision (%) |
| --- | --- | --- | --- | --- | --- |
| BitMapperBS (fast) | 957,664 | 956,665 | 999 | 95.77 | 99.90 |
| BitMapperBS (sensitive) | 958,362 | 957,082 | 1,280 | 95.84 | 99.87 |
| WALT | 901,278 | 898,890 | 2,388 | 90.13 | 99.74 |
| Bismark | 957,435 | 955,053 | 2,382 | 95.74 | 99.75 |
| BSMAP (gap) | 960,040 | 955,303 | 4,737 | 96.00 | 99.51 |
| BSMAP | 961,486 | 956,948 | 4,538 | 96.15 | 99.53 |
| GEMBS | 705,304 | 703,738 | 1,566 | 70.53 | 99.78 |
| ERNE-BS5 2 | NA | NA | NA | NA | NA |

ERNE-BS5 2 cannot be implemented successfully on this dataset. BSMAP denotes the results of BSMAP in default mode, which only supports mismatch alignment. By contrast, BSMAP (gap) denotes the results of BSMAP with gap alignment. Sensitivity is the percentage of reads that are aligned uniquely. Precision is the percentage of uniquely aligned reads that are reported correctly.

Although the exhaustive extension strategy improves the sensitivity and precision of BitMapperBS, it also results in more candidate locations that need to be extended. What’s worse, for three-letter WGBS aligners, there are more candidate locations than DNA sequence aligners. The reason is that in these WGBS aligners, the CT-converted seeds with low sequence complexity include a large number of occurrence positions on the reference genome. Thus, calculating the inexact alignment of these candidate locations is computationally expensive during extending step. To accelerate this time-consuming step, BitMapperBS adopts an efficient vectorized bit-vector algorithm [17], which extends multiple candidate locations simultaneously. In BitMapperBS, we modifiy this algorithm to support the asymmetric match between Cs and Ts directly. Besides, our implementation is developed based on AVX2 instructions to further improve its performance. It is able to calculate the inexact alignment of 8 candidate locations at once with up to 31 errors. Since current Illumina sequencing platforms usually produce WGBS reads ranging from 100bp to 300bp, it is enough to align them with up to 31 errors.

## 3 Results

To test the performance of BitMapperBS, we compared it with five state-of-the-art WGBS aligners, including Bismark [7], BSMAP [5], WALT [8], ERNE-BS5 2 [26] and GEMBS [27]. These aligners were run with default parameters unless otherwise stated, which are recommended by their authors (see Supplementary Section S2). Recent method BRAT-nova [9] was not evaluated in our experiments, since it cannot build index for the human genome successfully. This problem is also reported in [8]. In addition, BRAT-nova only supports single-threaded mode instead of multi-threaded mode, so that it cannot be as fast as other WGBS aligners on current multicore processors (see Supplementary Section S3). We also did not test another recent method BS-Seeker3 [11], since it was not run successfully for human genome on our experiments. Besides, BS-Seeker3 cannot significantly outperform existing aligners on widely used high quality WGBS datasets, and it is much more memory-consuming than other aligners (see Supplementary Section S3). The detailed discussion of BRAT-nova and BS-Seeker3 can be found in Supplementary Section S3. Other aligners like HPG-Methyl2 [28], BS-Seeker2 [10], BatMeth [29] and BWA-meth [30], were not tested due to their relatively poor performance presented in previous works [8, 31]. All experiments were performed using six CPU threads on a computer with a six-core Intel Core i7-8770k processor and 64 GB RAM, running Ubuntu 16.04. The indexes, reference genomes and reads were stored in a Solid State Drive (SSD) to minimize the loading time of WGBS aligners.

### 3.1 Performance comparison on simulated dataset

In this experiment, we focused on comparing the alignment sensitivities (the percentage of reads that are aligned uniquely) and precisions (the percentage of uniquely aligned reads that are reported correctly) of different methods. To this end, we generated 1 million simulated paired-end 150bp reads from human genome (GRCh38 assembly) using Sherman (http://www.bioinformatics.babraham.ac.uk/projects/sherman/) with an error rate of 1% and a bisulfite conversion rate of 98.5%. Sherman is designed specifically to simulate bisulfite sequencing reads, and has been widely used in previous studies [29, 31]. Since the original positions of simulated reads were known in advance, we could evaluate the precision of each WGBS aligner. In practice, to avoid incorrect alignment, most WGBS aligners only report the results of unique aligned reads. As shown in Table 1, all WGBS aligners presented comparable sensitivities except GEMBS. Although GEMBS achieved relatively high precision, the sensitivity of it was significantly lower than those of other aligners. GEMBS aligned 70.53% of reads uniquely, while most other aligners (BitMapperBS, Bismark and BSMAP) uniquely aligned more than 95% of reads. By contrast, BSMAP achieved the highest sensitivity among all methods, but its precision was the lowest. BitMapperBS shown the best precision, identified the most correctly aligned reads as well as the fewest incorrectly aligned reads, and achieved a good balance between sensitivity and precision.

### 3.2 Performance comparison on real datasets

To evaluate the running time (wall time) and the memory requirement of various WGBS aligners, we selected four large and real datasets with various read lengths produced by different high-throughput sequencing platforms. These four datasets were generated by first selecting the first 60 million paired-end reads from SRR948855 (100bp, HiSeq 2000), SRR6426282 (125bp, HiSeq 2500), SRR5453774 (151bp, NextSeq 500) and SRR6804872 (151bp, HiSeq X), respectively. After that, all reads were preprocessed by popular Trim Galore (https://www.bioinformatics.babraham.ac.uk/projects/trim_galore/) to perform the quality and adapter trimming. Following adapter and quality trimming, there were 59.6 million, 59.6 million, 59.2 million and 59.9 million paired-end reads retained on SRR948855, SRR6426282, SRR5453774 and SRR6804872, respectively. Note that unlike the simulated dataset, we cannot evaluate the precision of each WGBS aligner on real datasets without the original positions of reads. Thus, to evaluate the precision approximately, we measured the concordance of alignment results between the most widely used WGBS aligner Bismark and other aligners on four small real datasets. Each of these datasets was generated by selecting the first 1 million paired-end reads from its corresponding large dataset above.

As shown in Table 2, BitMapperBS was the fastest one among all WGBS aligners. Compared with the second fastest aligner GEMBS, BitMapperBS was about 9.0 times faster, 8.8 times faster, 12.3 times faster, and 9.7 times faster on SRR948855, SRR6426282, SRR5453774 and SRR6804872, respectively. In addition, BitMapperBS was much more sensitive than GEMBS on all datasets, and the memory requirement of GEMBS was 4.8 times larger than that of BitMapperBS. We also compared BitMapperBS with two widely used aligners BSMAP and Bismark. BitMapperBS was up to 85.7 times and 72.0 times faster than BSMAP and Bismark, respectively, and presented greater sensitivity. Note that although BSMAP (gap) and ERNE-BS5 2 achieved slightly higher sensitivities than BitMapperBS in some cases, the precisions of them were much lower than that of BitMapperBS (see Table 1 and Table S2 in Supplementary Section S4). For example, when aligning the first 1 million paired-end reads from SRR5453774, at least 99.30% of reads identified by Bismark were also aligned by BitMapperBS at same locations (see Table S2 in Supplementary Section S4). By contrast, only 97.09% and 92.03% of reads identified by Bismark were aligned concordantly by BSMAP (gap) and ERNE-BS5 2, respectively (see Table S2 in Supplementary Section S4). Besides, Table S2 in Supplementary Section S4 demonstrates that the alignment result of BitMapperBS was most concordant with that of Bismark. Actually, Bismark is one of the most widely used WGBS aligners so far. Another interesting finding is that BitMapperBS was scalable with the length of reads, while the performance of other WGBS aligners decreased when processing longer reads (see Table 2). Therefore, BitMapperBS is very suited for longer and longer WGBS reads produced by new high-throughput sequencing technologies in the near future. We also tested the performance of BitMapperBS with different numbers of CPU threads. As shown in Figure S1 in Supplementary Section S6, BitMapperBS presented good scalability with the increase of CPU threads.

**Table 2.** Performance of different aligners on real datasets from human genome.

| Dataset | Aligner | Thread | Running time<br>(wall time) (s) | Sensitivity<br>(%) | Memory usage<br>(GB) |
| --- | --- | --- | --- | --- | --- |
| SRR948855<br>(100bp, HiSeq 2000) | BitMapperBS (fast) | 6 | 186 | 87.3 | 6.8 |
|  | BitMapperBS (sensitive) | 6 | 206 | 87.8 | 6.8 |
|  | WALT | 6 | 10,630 | 80.3 | 16.9 |
|  | Bismark | 6 | 13,398 | 86.1 | 9.7 |
|  | BSMAP (gap) | 6 | 15,802 | 88.3 | 8.3 |
|  | BSMAP | 6 | 7,650 | 84.9 | 8.3 |
|  | ERNE-BS5 2 | 6 | 10,944 | 84.7 | 3.8 |
|  | GEMBS | 6 | 1,673 | 83.1 | 32.7 |
| SRR6426282<br>(125bp, HiSeq 2500) | BitMapperBS (fast) | 6 | 251 | 84.2 | 6.8 |
|  | BitMapperBS (sensitive) | 6 | 265 | 84.6 | 6.8 |
|  | WALT | 6 | 11,217 | 72.0 | 17.0 |
|  | Bismark | 6 | 14,936 | 79.7 | 9.7 |
|  | BSMAP (gap) | 6 | 30,349 | 84.5 | 8.3 |
|  | BSMAP | 6 | 12,856 | 78.8 | 8.3 |
|  | ERNE-BS5 2 | 6 | 10,603 | 84.2 | 3.8 |
|  | GEMBS | 6 | 2,219 | 61.5 | 32.8 |
| SRR5453774<br>(151bp, NextSeq 500) | BitMapperBS (fast) | 6 | 230 | 85.1 | 6.8 |
|  | BitMapperBS (sensitive) | 6 | 256 | 85.3 | 6.8 |
|  | WALT | 6 | 11,975 | 74.6 | 17.2 |
|  | Bismark | 6 | 15,937 | 80.8 | 9.8 |
|  | BSMAP (gap) | 6 | 61,189 | 83.9 | 8.4 |
|  | BSMAP | 6 | 19,712 | 79.9 | 8.4 |
|  | ERNE-BS5 2 | 6 | 13,698 | 85.5 | 3.8 |
|  | GEMBS | 6 | 2,831 | 45.8 | 33.0 |
| SRR6804872<br>(151bp, HiSeq X) | BitMapperBS (fast) | 6 | 298 | 84.9 | 6.8 |
|  | BitMapperBS (sensitive) | 6 | 303 | 85.1 | 6.8 |
|  | WALT | 6 | 11,007 | 71.4 | 17.2 |
|  | Bismark | 6 | 15,499 | 79.1 | 9.8 |
|  | BSMAP (gap) | 6 | 51,667 | 85.3 | 8.4 |
|  | BSMAP | 6 | 19,944 | 79.9 | 8.4 |
|  | ERNE-BS5 2 | 6 | 14,752 | 85.5 | 3.8 |
|  | GEMBS | 6 | 2,889 | 46.7 | 33.0 |

Running time (wall time), memory usage and sensitivity of each method when aligning the first 60 million paired-end reads of each dataset. Following adapter and quality trimming, there were 59.6 million, 59.6 million, 59.2 million and 59.9 million paired-end reads retained on SRR948855, SRR6426282, SRR5453774 and SRR6804872, respectively.

### 3.3 Performance of aligners with uniform error threshold

Please note that in the above experiments, the default error thresholds of different WGBS aligners are not uniform. For each aligner, its sensitivity is mainly affected by the error threshold. To compare the sensitivities of all aligners fairly, we tested them with uniform error threshold (about 4% of read length). Actually, about 4% of read length is the default error threshold of the popular WGBS aligner Bismark. The detailed configurations of different aligners in this experiment can be found in Table S3 in Supplementary Section S5. As shown in Table S5 in Supplementary Section S5, BitMapperBS was still one order of magnitude faster than other methods, and was one of the most sensitive WGBS aligners. Besides, BitMapperBS presented the best precision in this experiment. Like the results in Table 1 and Table S2 in Supplementary Section S4, the results in Table S4 and Table S6 in Supplementary Section S5 show that BitMapperBS identified the most correctly aligned reads on simulated dataset, and the alignment result of BitMapperBS on real datasets was most concordant with that of Bismark.

## 4 Conclusion

Due to the low sequence complexity and the large volume of the whole-genome bisulfite sequencing reads, it is difficult to align them onto the large reference genome accurately and efficiently. To solve this problem, we present an ultrafast WGBS aligner BitMapperBS. A novel version of FM-index, called three-letter FM-index, is proposed to accelerate the seeding step of BitMapperBS. In addition, BitMapperBS adopts the vectorized bit-vector algorithm to extend the candidate locations of each read efficiently. We also develop several other strategies specifically for the problems that are unique to the WGBS aligners. Experiments on both simulated and real datasets show that BitMapperBS is at most more than 70 times faster than popular WGBS aligners BSMAP and Bismark, and presents similar or greater sensitivity and precision. Although some recent WGBS aligners are faster than BSMAP and Bismark, BitMapperBS is still one order of magnitude faster than them, and achieves much better sensitivity.

## Acknowledgements

This work was partially supported by the National Nature Science Foundation of China under the grant No. 61672480 and the Fund for Foreign Scholars in University Research and Teaching Programs (B07033).

